# Unsupervised prioritization of gene-regulatory variants with isolation forests

**DOI:** 10.64898/2026.10.29.762447

**Authors:** Andreas W. Schmidt, Jonas Rosendahl, Helmut Laumen, Stephan Ossowski

**Author notes:** These authors contributed equally and share the last authorship (Helmut Laumen, Stephan Ossowski).

## Abstract

Most common variants occur in non-coding genomic regions. Numerous expression quantitative trait loci (eQTLs) can influence nearby gene expression. Across diverse traits and diseases, many genome-wide association study (GWAS) loci have been shown to colocalize with eQTLs, supporting gene-regulatory causal mechanisms. However, linkage disequilibrium (LD) obscures the identification of the causal variant(s), necessitating prioritization by functional annotations. Previously published variant prioritization tools relied on the machine-learning-based combination of diverse feature types like evolutionary conservation and chromatin information, whereas more recent deep learning tools learn and describe gene-regulatory DNA sequence grammar. However, the latter approaches often generate a large number of scores per variant. We designed a novel unsupervised machine-learning approach based on isolation forests to integrate the predictions from the broadly used Enformer deep learning model into a single gene-regulatory predictive score for each common variant. Our novel approach eQTL prediction with isolation forests (eQIF) demonstrates strong performance in predicting eQTLs and gene-regulatory GWAS variants. We show that top eQIF variants are located in evolutionarily conserved transcription factor binding site (TFBS) footprints, and that the frequency of top eQIF variants within overrepresented TFBS motifs recapitulates the motif information content. eQIF predictions are precomputed for 9,421,183 common variants and publicly available.

Graphical Abstract.
eQIF is a novel score for predicting the gene-regulatory impact of variants. eQIF is based on the top 20 principal components (PCs) from the publicly available precomputed Enformer deep-learning chromatin predictions overlapping common gnomAD variants. The PCs were provided to an isolation forest to learn global density within the multi-dimensional feature space. After training the model, it was applied to predict a global anomaly score (eQIF) for all gnomAD variants. eQIF was finally used for fine-mapped variant classification, functional variant discovery and precise prioritization of highly informative TFBS positions. Created in BioRender. Schmidt, A. W. (2026) https://BioRender.com/6dgiji3

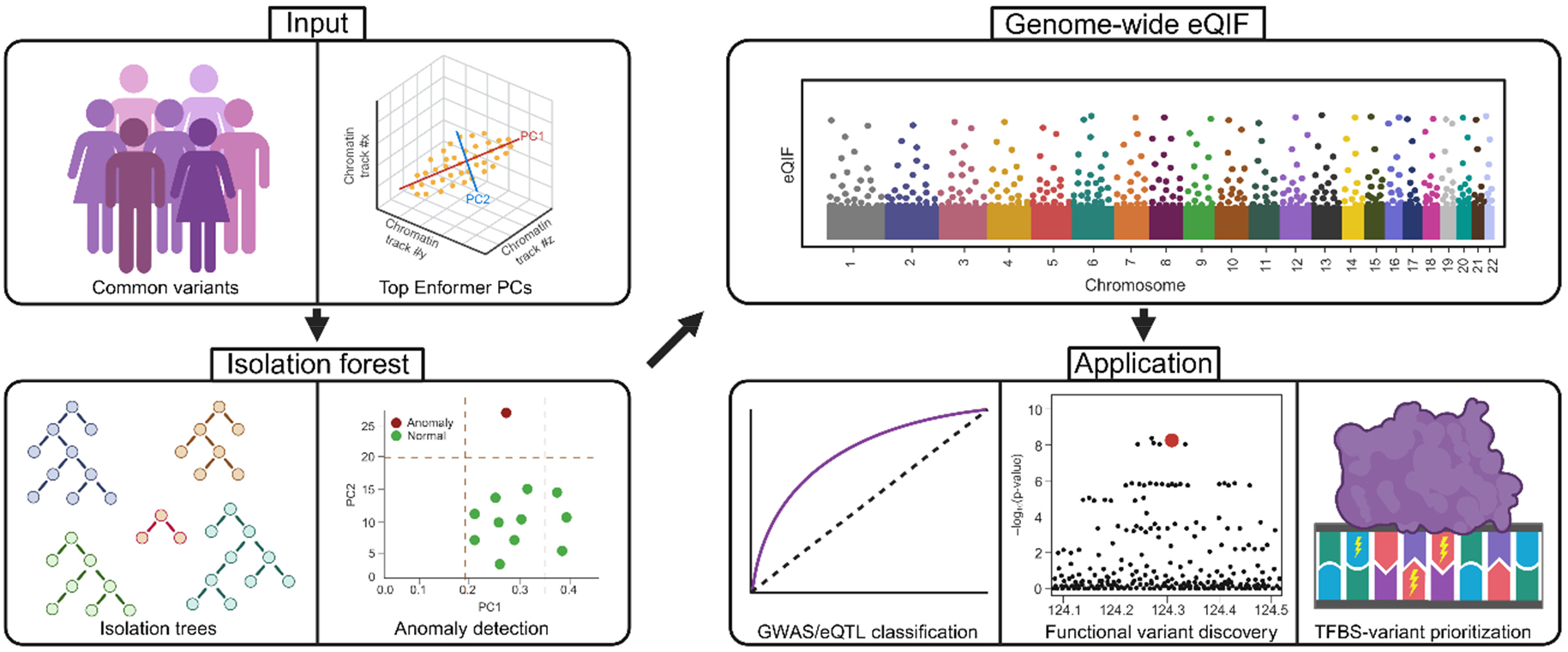

## Introduction

Approximately 10 million common variants of the human genome have a minor allele frequency > 0.5% in the European non-Finnish whole genome sequencing (WGS) cohort of gnomAD (1). These single nucleotide polymorphisms (SNPs) are mostly distributed in the non-coding regions of the genome and can influence the expression of nearby genes. A large number of expression quantitative trait loci (eQTLs) in multiple tissues have been mapped by the “Genotype-Tissue Expression” GTEx consortium (2). Previous genome-wide association studies (GWAS) coupled with eQTL colocalization (3) indicated gene-regulatory causal effects for quantitative traits (4), polygenic diseases (5, 6), and cancer (7). Closely located common variants can be co-inherited to the next generation during meiosis, leading to LD-blocks (8). The high correlation of alleles within LD-blocks impairs the discovery of causal variants, since the enclosed variants often show similar association levels. Advanced probabilistic data-driven fine-mapping methods can help to prioritize a smaller credible set of variants at a respective locus (9, 10). However, multiple variants might remain within the credible sets for eQTL or GWAS studies with small sample sizes or strong variant linkage. Furthermore, fine-mapping remains challenged in discovering multiple truly causal variants at a respective locus when two or more of them show high correlation.

Diverse variant impact prediction tools have been developed, which can help to prioritize causal variants and narrow down the experimental effort for validation, based on the integration of different molecular features. Combined Annotation Dependent Depletion (CADD) (11) and the recently proposed AlphaGenome Variant Impact (AVI) score (12) combine distinct features. They range from agnostic evolutionary conservation metrics (13, 14), missense variant effect scores from deep learning models (11, 15), to gene-regulatory chromatin annotations from large databases like Roadmap (16) or ENCODE (17) by supervised machine-learning to predict mechanism-agnostic variant impact. Though the mechanism-agnostic approach provides high flexibility, models like LINSIGHT (18) and DVAR (19), which integrate similar sets of features like CADD, but are specifically designed for gene-regulatory prediction, demonstrated stronger performances in predicting eQTLs and massively parallel reporter assay (MPRA) data (19). A significant strength of these models is that precomputed genome-wide variant effect predictions are available, enabling easy usage for variant prioritization.

In parallel, gene-regulatory-based deep learning models have emerged, which learn to directly predict a variant’s effect on chromatin features from DNA sequence grammar. These models showed largely improved representation of gene-regulatory variant effects (20, 21). One of the first available gene-regulatory deep learning tools was DeepSEA, which is based on a convolutional neural network (CNN) architecture and trained to predict a comprehensive dataset of chromatin annotations (22). More recently, the Sei framework expanded this model, by integrating more biological training data, by providing a reduced representation of variant effects in biologically meaningful sequence classes, and finally, even a single prioritization score per variant (23). The underlying CNN architecture of these types of models allows for learning local sequence architecture, but does not include distal enhancer-promoter interactions. This lack of biological information has been improved in the recently published Enformer model (20), which combines the CNN with a subsequent transformer architecture modeling distal sequence dependencies to predict chromatin features like histone modification levels, chromatin accessibility, and transcription factor binding. For each of those tracks, Enformer predicts the change of levels introduced by a genomic variant (delta score). In total, Enformer predicts more than 5,000 variant-level delta scores, representing all different molecular tracks across distinct tissues and cell types used for training. Those predictions and their projection to the top 20 principal components, which show similar predictive performance as the entire dataset, have been released as public resource (20). However, neither the full dataset nor the 20 linear projections of Enformer readily facilitate an easy manual genome-wide prioritization of variants, for which a single interpretable score is desirable.

Here, we integrated Enformer PCs to prioritize gene-regulatory variants across the genome. As a basic hypothesis for our work, we postulate that most variants are neutral and have no gene-regulatory effect, whereas a small fraction of variants are expected to be gene-regulatory and display rare Enformer patterns. To test this hypothesis, we chose the isolation forest (24) as the underlying unsupervised classification framework. This multi-dimensional outlier detection model splits the data based on the provided features and learns which data points are more isolatable than others. Strong data anomalies reflect rare genomic patterns, which might indicate functional effects. By using an unsupervised approach, we avoided potential tissue-/ or disease biases from training cohorts. Our “eQTL prediction with isolation forests” (eQIF) model outperformed the similarly designed unified Sei score and other previous models for global eQTL and gene-regulatory GWAS fine-mapped variant classification. At the prominent *SORT1* GWAS locus associated with LDL-C levels and myocardial infarction, as well as the novel melanoma risk locus *MX2*, eQIF was capable of prioritizing the respective experimentally validated risk variants (25, 26).

Finally, we extensively evaluate the properties of eQIF by regional gene-regulatory annotation (27), transcription factor, and evolutionary data. Our IF-based model eQIF runs computationally fast and allows for genome-wide precomputation of gene-regulatory variant effects.

## Methods

### Training an Enformer-PC-based isolation forest

To construct the input dataset for our novel model “eQTL prediction with isolation forests” (eQIF), we first defined a set of common human variants and annotated them with Enformer-derived features. Enformer-annotated variants were downloaded from https://console.cloud.google.com/storage/browser/dm-enformer/variant-scores/1000-genomes/enformer/PC20;tab=objects?prefix=&forceOnObjectsSortingFiltering=false. This dataset comprises variants from the 1000 Genomes Project with an allele frequency ≥ 0.5% and provides 20 principal components (PCs) derived from Enformer variant-effect predictions (20). As the Enformer annotations were provided in hg19 coordinates, variant positions were converted to the hg38 reference genome using the liftOver function from the R package rtracklayer (28). SNPs were extracted from the gnomAD v. 3.1.2 whole-genome sequencing (WGS) non-Finnish European (NFE) dataset and filtered for an allele frequency ≥ 0.5% and a “PASS” annotation in the quality column (1). In addition, the HLA-MHC region (chr6:25,000,000–34,000,000) was excluded because of its exceptionally high level of genetic variation compared with the remainder of the genome, which could bias model training. Only variants present in both the filtered gnomAD dataset and the lifted Enformer dataset were retained for all subsequent analyses. Following coordinate conversion, allele and strand inconsistencies were corrected based on the corresponding gnomAD annotations. The resulting 20 Enformer PCs were used as input features for eQIF.

For anomaly detection, we used isolation forest models as implemented in the R package isotree (29). For hyperparameter optimization, we performed five repeated random 80:20 training-test splits of the complete genome-wide variant dataset (Monte Carlo cross-validation). In each iteration, isolation forest models were trained on a randomly selected subset comprising 80% of all variants. Model performance was subsequently evaluated on the remaining 20% of variants, restricted to those overlapping the predefined eQTL case and control cohorts. The random splits were generated independently using different random seeds, allowing variants to appear in multiple test sets across iterations. The optimal parameter combination was selected based on the mean AU-ROC across the five iterations. We evaluated sample_size, defining the number of genomic variants per tree; and ndim, which determines the number of features considered at each split. Values of 1,000 and 10,000 were tested for sample_size, and 1 or 2 for ndim, corresponding to a standard isolation forest or an extended isolation forest, respectively (30). Furthermore, we investigated feature weighting using the column_weights parameter by comparing models with equal feature weights to models in which the 20 Enformer PCs were assigned linearly decreasing weights according to their rank, ranging from 20 for PC1 to 1 for PC20. The number of trees was set to 500 in all models. All possible parameter combinations were assessed, resulting in 8 candidate models. The best model was then selected based on the mean AU-ROC computed with the R-package precrec (31).

Using the selected parameter set, the final eQIF model was subsequently trained in an unsupervised manner on the complete set of retained common variants, using their 20 Enformer PCs as input features. The trained isolation forest was then used to assign an anomaly score to each SNP in the dataset.

### Evaluation with fine-mapped eQTL, gene-regulatory GWAS and MPRA data

To evaluate eQIF across different regulatory and disease-associated variant settings, we assembled several validation cohorts. All validation cohorts were restricted to variants contained in both the gnomAD-derived variant set and the corresponding Enformer-annotated dataset used for eQIF model development.

Fine-mapped eQTL variants were extracted from the GTEx database (v. 8). The fine-mapping was originally conducted with the tool “causal variants identification in associated regions” (CAVIAR), which provides a posterior probability of fine-mapping for each variant to be a causal eQTL for each gene in each analyzed tissue (9). To obtain a set of highly credible causal eQTL variants, we filtered the entire GTEx dataset for a CAVIAR posterior probability ≥ 0.95. Control variants were defined by a maximum posterior probability ≤ 0.01 and were further matched by allele frequency (deviation of 0.1 * AF allowed) and the distance to the nearest basic transcript transcriptional start site (TSS) of a coding gene (± 250 bp). The underlying gene and transcript information was extracted from the GENCODE v. 44 dataset, and the allele frequency information required for matching was extracted from the gnomAD v. 3.1.2 database (1). In this manuscript, the obtained cohort is termed “eQTL cohort”. Furthermore, a subset from this eQTL-dataset was formed by keeping only variants remaining within GENCODE coding sequence regions for coding eQTL analysis, which we term “eQTL-coding cohort”.

Furthermore, a testing cohort of gene-regulatory trait-altering variants was defined to study eQIF’s predictive performance on eQTL variants influencing phenotypes. For this purpose, first fine-mapped GWAS variants were extracted from CAUSALdb2, which provides a large collection of fine-mapping results for > 10,000 traits (32). To define fine-mapped GWAS variants, all SNPs with a “Sum of Single Effects” (SuSiE) (10) fine-mapping probability ≥ 0.95 were extracted. Control variants were required to have a posterior probability ≤ 0.01 by SuSiE. Next, case variants were filtered for being both fine-mapped eQTLs, as defined before, and fine-mapped GWAS variants. Variants annotated as controls for both eQTL and GWAS were then matched by distance and allele frequency. Together, the case and control variants formed the “GWAS-eQTL cohort”.

Additionally, a massively parallel reporter assay (MPRA) cohort was taken directly from Siraj et al. (33). This dataset consists of 221,412 fine-mapped variants (78,238 trait-associated variants and 148,805 eQTL variants, including 5,631 variants shared between the two groups) and additionally 86,064 variants based on annotation or location matching or simulated null-trait associations. Case variants were defined on significant allelic differences in an MPRA assay in at least one of 5 cell types. Control variants were defined by the absence of MPRA differences in all cell types.

After defining the validation cohorts, ROC and precision-recall (PR) analysis were conducted to compare the novel eQIF score with other previously published models. For classification of the GWAS-eQTL and the MPRA cohort additional scores were formed and included by taking the rank product of either eQIF and AVI or eQIF and seqclass_max_absdiff, respectively. The scores were termed rp_EA and rp_ES respectively. The statistical significance of pairwise differences in ROC was evaluated with DeLong’s test implemented in the pROC R-package (34) and Benjamini-Hochberg (BH) correction. CADD v. 1.7 and AVI, which provide mechanism-agnostic variant prioritization, were included (11, 12). Furthermore, the hg19-based gene-regulatory DVAR (19) and LINSIGHT (18) scores were lifted to hg38 and also included in the comparisons.

Finally, the CNN-based Sei seqclass_max_absdiff score was included in the analysis. Like Enformer, Sei is a sequence-based gene-regulatory model, but it is founded on a CNN framework. It computes a large number of chromatin predictions for each variant, which it summarizes into 40 sequence classes, which are further reduced to a single unsupervised score, seqclass_max_absdiff (23).

Missing values for any model were imputed by the median.

### Computation of the seqclass_max_absdiff score

The unsupervised deep-learning-based gene-regulatory variant impact predictor seqclass_max_absdiff score from Sei was computed for all investigated variants. First, the VCF of common European SNPs of high quality was provided to Sei. Next, the 1_variant_effect_prediction.sh script, which computes chromatin feature levels for reference and alternative sequences, was run in GPU mode by adding the parameter --cuda. The final variant effect scores for each chromatin track were computed with the 2_varianteffect_sc_score.sh script. This step gives sequence class scores, which represent the differential variant impact from the reference and alternative sequence for specific biological annotations (sequence classes), e.g., promoter or CTCF-binding. From this table, we computed the seqclass_max_absdiff score for each variant by taking the maximum of the absolute sequence class scores.

### Prioritizing functionally validated risk variants

The variant *rs12740374*, which is experimentally validated to alter the expression of *SORT1*, leading to increased LDL-C levels and myocardial infarction, and the further strongly associated non-functional variants *rs11102967*, *rs660240*, *rs629301*, and *rs646776* were annotated with eQIF, CADD, LINSIGHT, DVAR, AVI, and Sei. Additionally, the recently MPRA-validated melanoma gene-regulatory risk variant *rs398206*, and the non-functional, high-LD variants *rs408825*, *rs364525*, *rs416981*, *rs438646*, and *rs443099* from the *MX2* locus with a Probabilistic Annotation INTegratOR (PAINTOR) (35) fine-mapping probability ≥ 5% were annotated. Next, the respective prediction scores were ranked across the experimentally investigated variants to match them on a comparable scale. Finally, the ranked scores were used to investigate each tool’s capacity to predict the causal variant on rank 1.

### Evolutionary analysis

All included SNPs were annotated with evolutionary information based on the phyloP scores from vertebrate, mammal, and primate alignments (36). Then all variants were categorized for each of the three scores into “conserved” and “non-conserved” if they were in the respective set of top 1% phyloP scores or not. These binary outcomes were then modeled by regressing the scaled eQIF score. After building the models, the effect sizes were extracted and transformed to odds ratios, and the respective 95% confidence intervals were computed.

### Gene-regulatory analysis

All SNPs were annotated with gene-regulatory regional information from the SCREEN database (27) and genome-wide DNase-seq-based TFBS footprint from Vierstra et al. (37). The SCREEN data were further subset into promoter and enhancer elements. The genome was then divided into non-regulatory, enhancer, enhancer footprint, conserved enhancer footprint, promoter, promoter footprint, and conserved promoter footprint regions. Conservation was given if a variant fell into the top 1% global conserved variants by the phyloP score from the mammal alignment. For illustration, boxplots were generated.

### TFBS analysis of top scoring eQIF variants

After obtaining the global eQIF score, the top 5,000 variants were extracted as a representative set of highly credible gene-regulatory candidates. Next, Analysis of Motif Enrichment (AME) was conducted on SNP-centric windows to check for enriched transcription factors at top eQIF variants compared to shuffled background sequences (38). The underlying database for TFBS motifs was the non-redundant JASPAR v. 2026 vertebrate collection (39). Since the maximum motif length within the database was 33 bp, the windows were designed by extending the top 5,000 SNP positions by 33 bp. Prior to AME analysis, low-complexity regions were removed by dust analysis implemented in the MEME Suite (40). The E-value threshold to report an enriched motif by AME was set to 0.05. After obtaining the list of significantly enriched motifs, a fold-change filter of 2 was applied to remove frequently occurring hits with low effect sizes. Since TFBS motifs of different TFs can highly correlate, the obtained motifs were further grouped by both binding domain similarity of the underlying transcription factor and sequence-based similarity. The underlying grouping information was extracted from the JASPAR database. Finally, based on its broad appearance in the scientific literature, *CTCF*, and the strongest enriched factor *ETS1* (see Results) were extracted and provided to the “Find Individual Motif Occurrences” (FIMO) tool using default parameters (41). This tool computes the presence and exact location of a TFBS within provided sequences. We applied FIMO to regions centered on the top 5,000 SNPs. The window size was determined precisely by extending each variant position in both directions by the length of the TFBS for *ETS1*, 9, and *CTCF*, 15, to exclude motifs not affected by the SNPs. Within the discovered motif positions of *CTCF* and *ETS1*, we annotated the frequency of the top 5,000 eQIF variants and linked it with the TFBS information content across the letter sequence by measuring Spearman correlation.

## Results

### eQIF – a novel unsupervised machine learning tool for gene-regulatory variant effect prediction

The transformer model Enformer (20) marks a milestone in understanding gene-regulatory biology, since it includes distal interactions based on its transformer architecture. However, the utility of Enformer for variant prioritization is limited due to the complexity of its 5,313 outputs. Thus, we aimed to merge the highly informative publicly available top 20 Enformer PCs into a single genome-wide gene-regulatory variant effect score for all SNPs to prioritize eQTLs. In order to avoid biases, we trained the unsupervised isolation forest (24) model “eQTL prediction with isolation forests” (eQIF) on the PCs for all European SNPs with AF > 0.5%. The IF learns global density within the multi-dimensional feature space and provides an anomaly score for each variant.

### eQIF optimization and global evaluation

For eQTL evaluation of the genome-wide eQIF scores, fine-mapped variants from the GTEx database were contrasted against location and allele-frequency-matched control variants with low fine-mapping probability. In the beginning, eQIF hyperparameters were tuned on the eQTL cohort. The resulting models were evaluated in their eQTL predictive performance by using variants overlapping the eQTL cohort. Overall, choosing 500 trees each consisting of 1,000 genomic variants, considering just a single variable per split, and weighting the top 20 PCs, provided the best results in terms of AU-ROC (Suppl. Fig 1A). Different parameter sets showed only marginal performance differences (Suppl. Fig 1A&B) in absolute terms, demonstrating the low parameter sensitivity and high robustness of eQIF models. We next computed eQIF with the optimized parameters on the entire set of all SNPs and compared it to the previously published scores CADD, LINSIGHT, DVAR, AVI, and Sei (11, 12, 18, 19, 23). The predictions from eQIF showed low global similarities with CADD and AVI, which both can predict multiple distinct molecular mechanisms in a single score (Suppl. Fig 2). Moderate levels of correlation were observed with the specific gene-regulatory prediction tools LINSIGHT, Sei, and DVAR (Suppl. Fig 2). Notably, eQIF showed the strongest eQTL classification performance (AU-ROC = 0.77, AU-PRC = 0.13), followed by DVAR, LINSIGHT, Sei, AVI, and CADD in terms of AU-ROC and followed by AVI, Sei, DVAR, LINSIGHT, and CADD in terms of AU-PRC (Figure 1A&B; Suppl. Table 1). Furthermore, eQIF strongly outperformed all competitors in terms of AU-ROC, which was highly significant after BH-correction (Suppl. Table 2). Next, we specifically investigated eQIF’s performance in prioritizing eQTLs within coding regions, an often neglected category, which currently gains rising interest in the scientific community (42).

**Figure 1:**
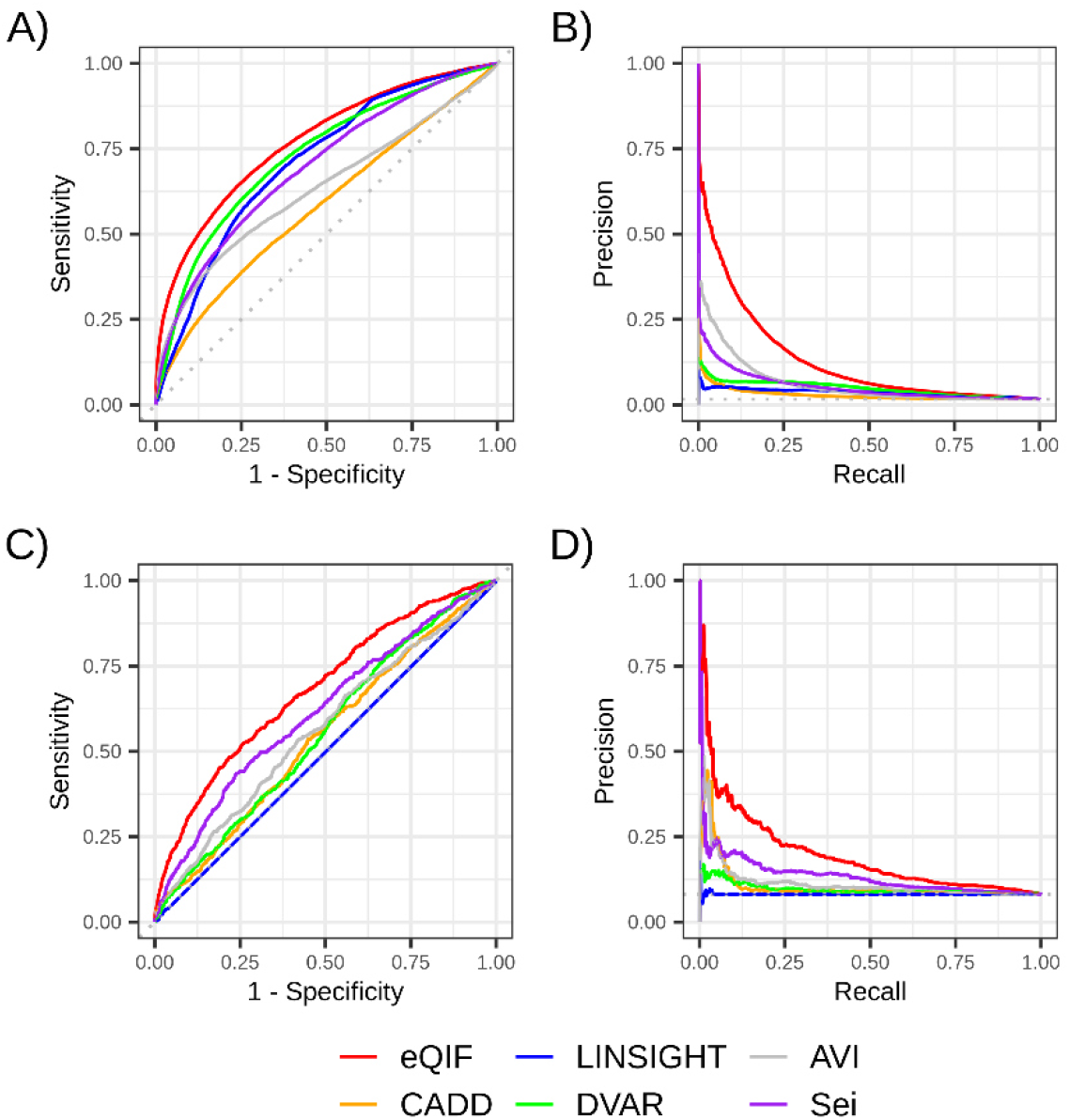
Comparing eQIF with previous tools for eQTL classification. A) ROC and B) PRC analysis for the eQTL cohort. eQTL classification was evaluated on a group of variants from the GTEx v.8 dataset with high fine-mapping probability versus a control dataset of matched variants with low fine-mapping probability. eQIF was compared to CADD v. 1.7, LINSIGHT, DVAR, AVI, and seqclass_max_absdiff from Sei. The number of case and control variants for the eQTL cohort was 12,727 and 767,512, respectively. C) ROC and D) PRC analysis for the eQTL-coding cohort. For coding eQTL classification variants were reduced to GENCODE coding regions of basic transcripts for coding genes. The number of case and control variants for the eQTL-coding cohort was 553 and 6,244, respectively.

Importantly, eQIF showed the strongest performance (AU-ROC = 0.68, AU-PRC = 0.2) of all tools on separating eQTL from control variants within coding regions (Figure 1C&D; Suppl. Table 1). Again, the ROC curve was significantly different compared to all other tools (Suppl Table 2). Intriguingly, the observed near-random performance for LINSIGHT within coding regions is mainly driven by data sparsity, since it is only computed for non-coding regions (18), thus overall reducing its utility in gene-regulatory variant prioritization.

### Gene-regulatory deep learning models show superior performance in MPRA prediction

After classification of global fine-mapped eQTLs, we next investigated the performance of eQIF in an *in vitro* massively parallel reporter assay dataset summarized from five cell lines (33). All variants showing allele-specific effects were contrasted with non-allele-specific variants. The highest AUC values were obtained for the seqclass_max_absdiff score from the Sei framework (AU-ROC = 0.68, AU-PRC = 0.28), closely followed by eQIF (AU-ROC = 0.67, AU-PRC = 0.27), both outperforming the remaining models showing weaker performances in their respective curves and AUC values (Suppl. Table 1 & Suppl. Fig 3). Notably, combining eQIF with the seqclass_max_absdiff score by a rank product further improved both AU-ROC and AU-PRC, suggesting that the two scores capture complementary information and mutually enhance predictive performance (AU-ROC = 0.69, AU-PRC = 0.30; Suppl. Fig 4). All pairwise ROC differences between all tools were significant by DeLong’s test after BH correction (Suppl. Table 2). Overall, the analyses of MPRA data indicate superior performance of purely deep learning-based gene-regulatory predictors for variant effect classification of exogenous reporter constructs.

### Prioritization of gene-regulatory GWAS variants

eQTLs represent one of multiple causal mechanisms at GWAS loci as indicated by numerous GWAS-eQTL colocalization studies (5–7). Challenging the high predictive performance of our eQIF model in eQTL prioritization, we next investigated the performance on predicting trait-altering gene-regulatory GWAS variants. Thus, we designed a second testing cohort based on case variants that were fine-mapped by both GWAS data from a large compendium of traits, including disease cohorts (32), and GTEx eQTLs (2). For this GWAS-eQTL cohort control variants were designed to be neither fine-mapped by GWAS nor eQTL analysis. AU-ROC revealed an overall higher performance by all predictors for GWAS-eQTL compared to the eQTL cohort (Suppl. Table 1 & Figure 2). eQIF showed the strongest performance as compared to all other precomputed scores in terms of AU-ROC (0.86) and was on par with AVI in terms of AU-PRC (0.23) (Figure 2A&B). Notably, eQIF considerably outperformed also the Sei seqclass_max_absdiff score (AU-ROC = 0.79, AU-PRC = 0.11), which is also a gene-regulatory deep learning-based variant representation, with a more than 2 times increase in AU-PRC. DeLong’s test indicated a significant outperformance of eQIF over its competitors after BH correction (Suppl. Table 2). When integrating eQIF with AVI by a rank product, a high increase in AU-PRC was observed, indicating that the two approaches provide complementary predictive information (AU-ROC = 0.87, AU-PRC = 0.33; Suppl. Fig 5).

**Figure 2:**
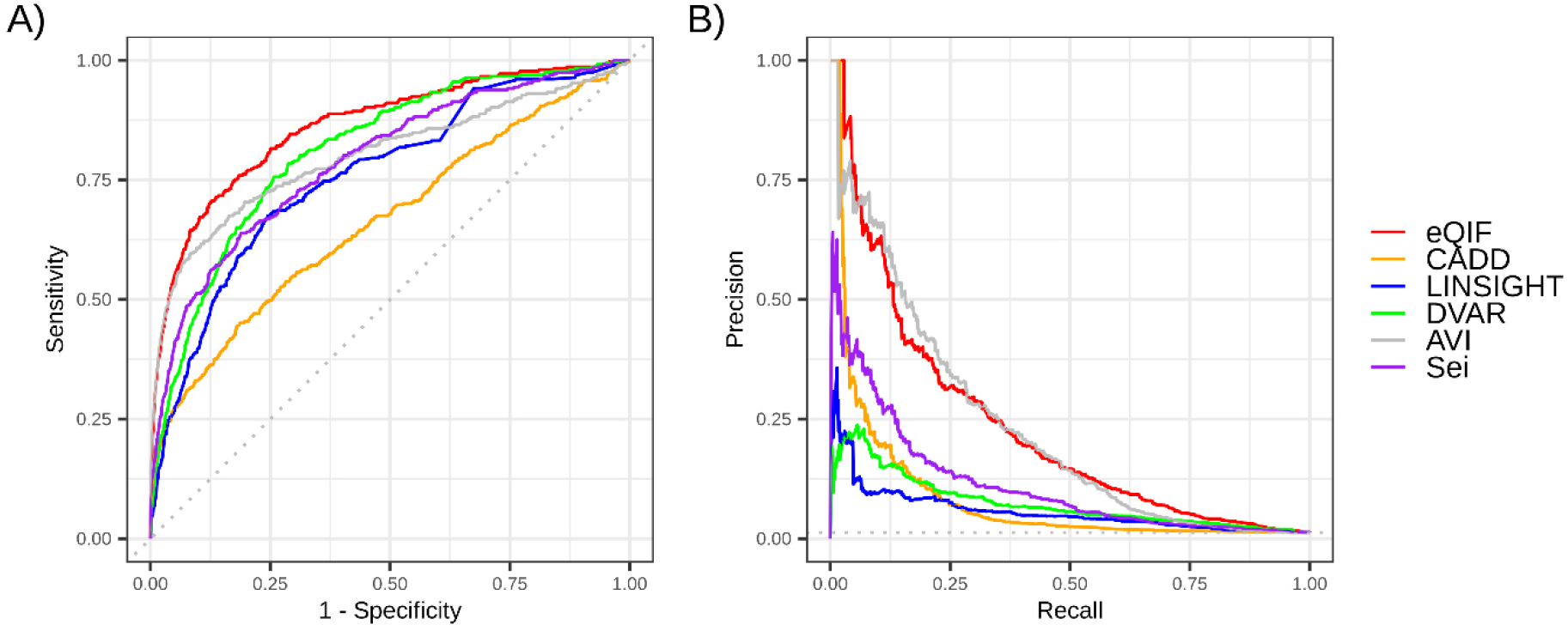
Comparing eQIF with previous tools for gene-regulatory GWAS classification. A) ROC and B) PRC analysis. GWAS-eQTL classification was evaluated on a positive group of variants strongly fine-mapped by both the CAUSALdb2 dataset and GTEx v.8 and a control dataset of variants with low fine-mapping probabilities. eQIF was compared to CADD v. 1.7, LINSIGHT, DVAR, AVI, and seqclass_max_absdiff from Sei. The number of case and control variants is 357 and 27,394, respectively.

### eQIF prioritizes functional causal variants at known GWAS risk loci

After investigating eQIF’s performance in gene-regulatory GWAS prediction with a genome-wide case-control setup, we next hypothesized that eQIF might also be suitable for prioritizing the causal gene-regulatory variant within GWAS haplotypes. To investigate this hypothesis, we selected the *SORT1* (25) and *MX2* (26) loci associated with both plasma low-density lipoprotein cholesterol (LDL-C) and myocardial infarction and melanoma, respectively. Both loci can be considered bona fide loci, since the causal variants have been discovered by experimental validation. For *SORT1*, the *rs12740374* variant has been shown to act as a *SORT1* enhancer variant by luciferase experiments. In total, we picked 5 variants from Musunuru et al. (25), which were all highly associated with LDL-C levels and myocardial infarction. At the *MX2* locus, the annotation-informed fine-mapping approach PAINTOR provided posterior probabilities ≥ 5% for six candidate risk variants (26). Subsequent MPRA analysis finally revealed *rs398206* to upregulate *MX2* expression levels (26). We annotated the extracted variants with ranked eQIF, CADD, LINSIGHT, DVAR, AVI, and Sei scores for both loci. At *SORT1*, eQIF, AVI, and Sei prioritized *rs12740374* on rank 1, whereas the remaining tools only prioritized it on rank 4. At the *MX2* locus, eQIF, AVI, Sei, and DVAR prioritized *rs398206* on rank 1, whereas LINSIGHT and CADD found it on ranks 2 and 3, respectively (Figure 3). Notably, eQIF scores for *rs12740374* and *rs398206* were 0.80 and 0.78, representing the top 0.04% and 0.08% strongest predicted variants across the genome, respectively. Both results indicate the genome-wide strong prioritization of our novel approach for both variants. Sei and AVI also showed a strong global prioritization of *rs12740374* and *rs398206*, whereas the remaining tools showed weaker performance (Suppl. Table 3).

**Figure 3:**
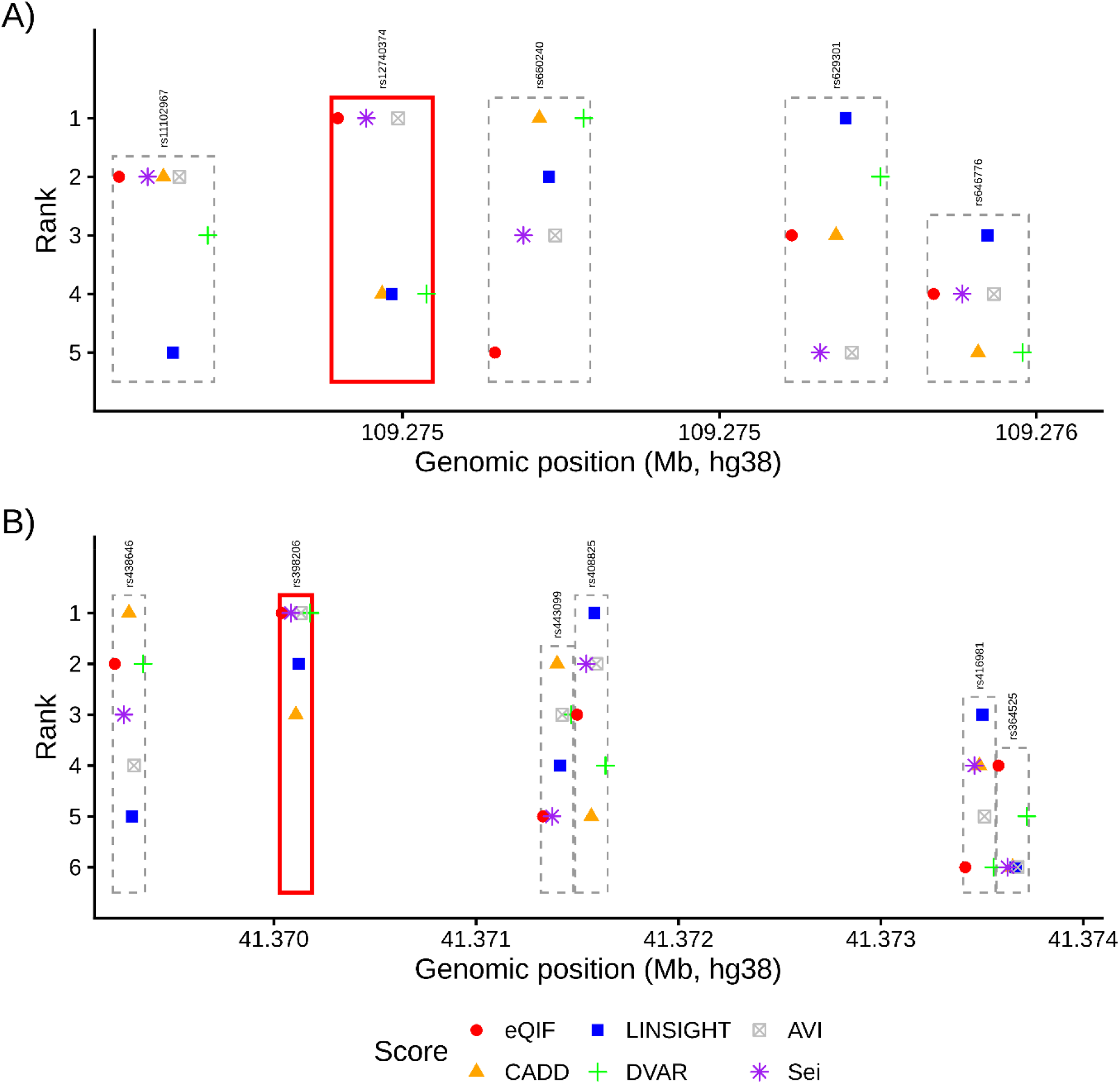
Prioritization of variants at experimentally evaluated GWAS loci. A) Five variants which were previously experimentally investigated at the SORT1 risk haplotype were annotated with the variant impact scores CADD, LINSIGHT, DVAR, AVI, Sei and our novel eQIF. The different tool predictions were ranked (rank 1-5) across the 5 variants for comparison on a similar scale. Rectangles indicate the distinct SNP positions. The causal SNP rs12740374 is marked in red. B) Six variants previously experimentally investigated at the MX2 risk haplotype were annotated with the variant impact scores CADD, LINSIGHT, DVAR, AVI, seqclass_max_absdiff (Sei) and our novel score eQIF. The different tool predictions were ranked (rank 1-6) across the 6 variants for comparison on a similar scale. Rectangles indicate the distinct SNP positions. The causal SNP rs398206 is marked in red. For each SNP position the tools are slightly horizontally shifted to avoid overlap in case of similar ranking.

### Biological characterization of eQIF predictions

After evaluating the predictive performance of eQIF for gene-regulatory variants, we evaluated its biological characteristics. First, we investigated whether eQIF is predictive of evolutionary conservation. We grouped all SNPs into conserved and neutral sites by primate, mammal, and vertebrate phyloP scores (14, 36). We discovered significant positive associations of eQIF with all three conservation scores, showing a 1.3-1.39-fold increase in the odds of a variant being conserved for one standard deviation increase in eQIF (Figure 4A, Suppl. Table 4). Notably, the enrichments were similar across the different groups of species, with the highest levels reached for the mammal phyloP. Thus, variants prioritized by eQIF tend to be located in evolutionary conserved regions and might have higher fitness consequences compared to low eQIF variants.

**Figure 4:**
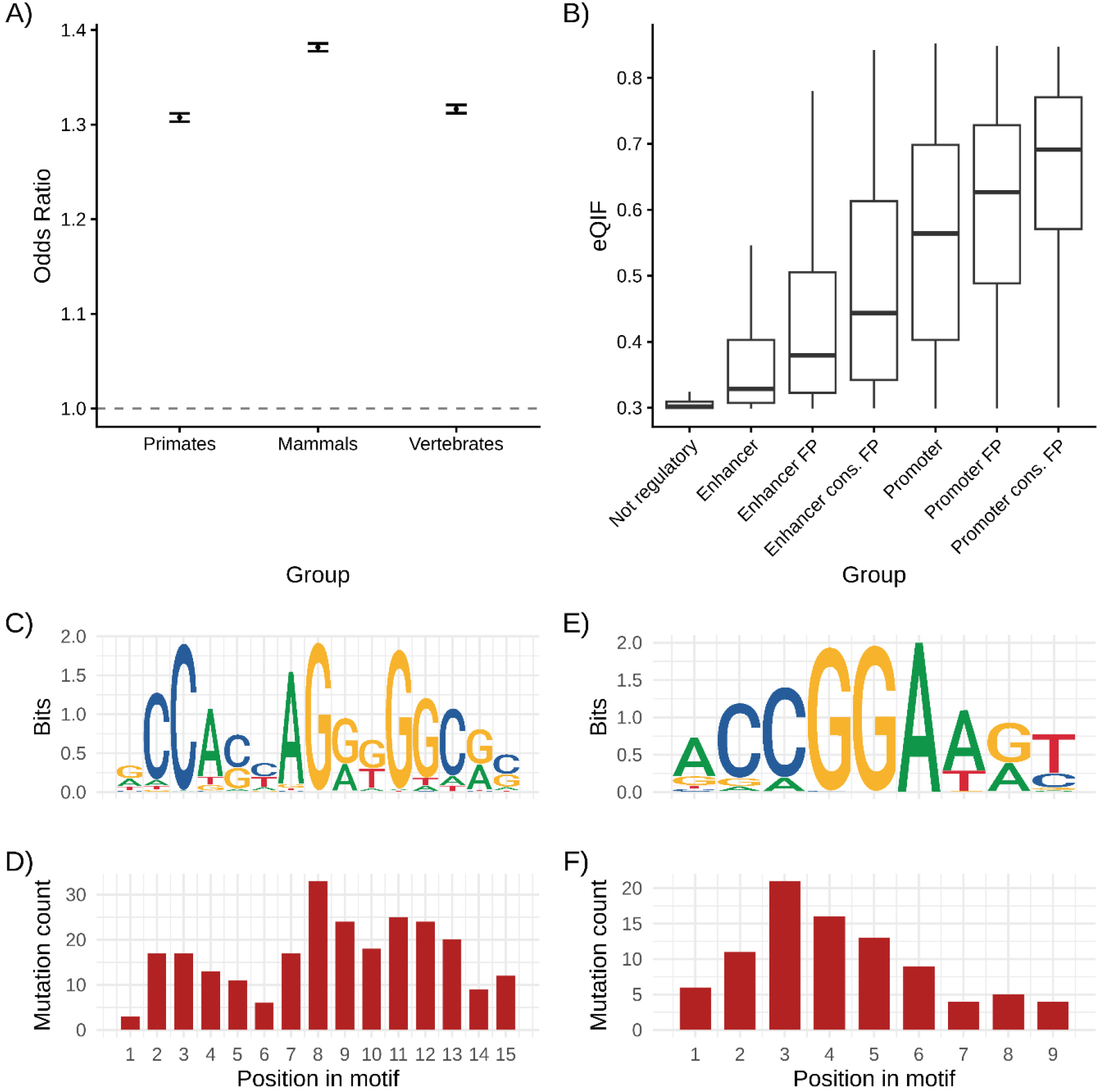
Biological characterization of eQIF. A) Global eQIF scores were regressed against the binary status of a variant to be conserved or neutral. Conservation was defined if a variant was in the top 1% of phyloP scores for primates, mammals or vertebrates, respectively. Odds ratios and 95% confidence intervals are shown. B) All SNPs were grouped by non-gene-regulatory, promoter and enhancer annotation. Promoter and enhancer positions were further subset by overlap with a TFBS footprint (FP) and a high level of mammal conservation (cons.). eQIF scores are shown across the different gene-regulatory annotations. C) CTCF motif (MA0139.2, JASPAR database). D) The number of SNPs located at each CTCF motif letter across the top 5,000 eQIF SNPs. E) ETS1 motif (MA0098.4), JASPAR database. F) The number of SNPs affecting each ETS1 motif letter across the top 5,000 eQIF SNPs. The information content at motifs is measured in Bits, ranging from 0 to 2.

When investigating the distribution of eQIF scores in different gene-regulatory genomic regions, we found the lowest scores for variants not located in any gene-regulatory element, while enhancer and promoter variants exhibited progressively higher eQIF scores. The enrichment of higher eQIF scores within promoter and enhancer elements was further improved when considering only TFBS-footprint regions and even stronger when filtering the positions for high evolutionary conservation levels (Figure 4B).

Given that our gene-regulatory region analysis indicates high-scoring eQIF variants to be enriched within transcription factor footprints, we next investigated which TFBS are enriched with the 5,000 genome-wide top-scoring eQIF variants. We discovered 142 significantly enriched TFBS with high fold-change (>2), which could be categorized into 11 and 23 groups based on binding domain and sequence-similarity clustering, respectively (Suppl. Figure 6). The strongest enrichment among all TFBS was found for *ETS1* (E-value = 1.36 x 10^-92^) belonging to the class of tryptophan cluster transcription factors (Suppl. Table 5 & Suppl. Fig 6). This second largest TF family in our dataset is characterized by evolutionary conserved hydrophobic tryptophans at their DNA binding sites (43). *ETS1* and other top-enriched ETS/ETV factors within our analysis are well-known drivers of cellular development and carcinogenesis (44, 45). Furthermore, 13 of the significant TFBS were the only instances for their respective annotated sequence-based classes, allowing to identify the underlying transcription factor. Among those factors, we discovered *CTCF* (E-value = 3.34 x 10^-15^), a well-studied ubiquitously expressed protein playing a key role in organizing chromatin architecture (46). *CTCF* is broadly recognized as master regulator of genome topology, with its binding sites being highly evolutionary conserved compared to other transcription factors (47).

Finally, we tested if eQIF scores resemble the information content at each position of the enriched motifs (represented by the position weight matrix). For *CTCF* and *ETS1*, we observed strong levels of positive Spearman correlation (*CTCF* ρ = 0.70, p-value = 0.004; *ETS1* ρ = 0.67, p-value = 0.049), respectively, between the eQIF top-5000 variant count and the weight matrix position information content. Thus, top eQIF scores within the TFBS motifs of *CTCF* and *ETS1* are predominantly found at highly informative motif positions (Figure 4C-F).

## Discussion

We present eQIF, a novel gene-regulatory variant impact score, which is unsupervised by design and genome-wide precomputed for all SNPs with AF ≥ 0.5% in the European population. The score is based on the multi-dimensional unsupervised outlier detection tool isolation forest (IF) (24) and integrates the highly informative top principal components (PCs) from the precomputed Enformer dataset (20). Enformer predictions summarize gene-regulatory variant effects from a large number of tissues, replicates and different molecular assays by using a sequence-based deep learning model. Specifically, Enformer, which is founded on a transformer model, also captures distant interactions such as enhancer-promoter dependencies (20). Our model eQIF integrates this high complexity into a single unsupervised score based on the simple assumption that gene-regulatory variants should be rare across the genome, and so should be their feature patterns.

We demonstrated that eQIF shows superior performance over the previous predictors CADD, LINSIGHT, DVAR, AVI and Sei in global eQTL classification within the large fine-mapped eQTL cohort (2). One major application in the future, based on eQIF’s strong performance, is candidate variant prioritization after eQTL fine-mapping within remaining credible sets, which will guide experiments to improve both our understanding of gene-regulatory biology and the discovery of causal variants.

Notably, we found eQIF to clearly outperform the other scores when classifying eQTLs within coding regions, followed by the Sei, the second gene-regulatory deep learning-based score. The strong/superior performance of both models may be explained by their exclusive gene-regulatory focus, whereas other scores might get confounded by inclusion of conservation and molecular mechanisms within coding sequences. Although enhancer activity within coding exons was already reported in the 1990s (48), most exon sequencing studies focused on missense-related mechanisms. However, this overlooked class of coding enhancer variants has been shown to play a particular role in gene regulation and distinct cancer types (42).

eQIF’s strong prediction of gene-regulatory variants is furthermore supported by our exogenous MPRA classification analysis on a large number of variants. eQIF was outperformed only by Sei, potentially reflecting an advantage on artificial reporter assay data, where the broader genomic context modeled by Enformer in the eQIF scores is not present. Lentiviral MPRA data, owing to their integration into the genome, have been shown to better recapitulate *in vivo* genomic chromatin annotations (49). Future large-scale endogenous lentiviral MPRA atlases could be used to investigate eQIF’s predictive performance in a more biologically plausible setting.

Moreover, we demonstrated eQIF’s superior performance in the prediction of fine-mapped GWAS variants (32), which were also annotated to be fine-mapped eQTLs in the GTEx dataset (2). Notably, eQIF again outperformed the recently published seqclass_max_absdiff score from the Sei framework, which is similar to eQIF by its design as unsupervised representation from a large sequence-based chromatin-effect model (23). One explanation for the outperformance of eQIF may be that Sei is a CNN architecture not including distant gene-regulatory interactions, whereas the transformer-based Enformer also captures such distant interactions.

Besides genome-wide classification of variants, we also investigated eQIF’s performance in fine-mapping by analyzing the well-characterized *SORT1* and *MX2* GWAS loci (25, 26). At the *SORT1* locus eQIF, Sei and AVI scores prioritized the experimentally validated gene-regulatory risk variant *rs12740374*, whereas CADD, LINSIGHT and DVAR failed. At the *MX2* locus, previously published annotation-based fine-mapping with PAINTOR revealed the highest posterior probability for *rs364525*. Subsequent MPRA analysis, however, revealed *rs398206* to be the gene-regulatory causal variant. eQIF, Sei, AVI and DVAR were capable of prioritizing *rs398206*, whereas the frequently used tools CADD and LINSIGHT did not succeed. Taken together, our GWAS results indicate eQIF’s strong performance in classifying fine-mapped GWAS and eQTL variants. Thus, one potential application of eQIF is to support fine-mapping following eQTL colocalization analysis (3) in future studies. This could be particularly useful when multiple variants remain in the final credible set.

### Limitations

Overall, eQIF provides a better global ranking of gene-regulatory GWAS variants compared to AlphaGenome Variant Impact (AVI), as indicated by AU-ROC. However, for the top predicted variants, both tools showed equal performance, as indicated by AU-PRC. One reason for the competitive performance of AVI for top predicted variants might be its supervised training procedure to learn global deleteriousness by using allele frequency as a proxy and using conservation measurements as part of the feature set. Overall deleteriousness might play a bigger role in gene-regulatory GWAS variants than pure eQTLs, since a significant fraction of GWAS loci is associated with disease traits (32). Furthermore, AVI includes AlphaGenome annotations, which span a larger context window than Enformer, thus integrating more distant eQTL effects (50). Theoretically, eQIF could be easily improved by running the IF on AlphaGenome annotations; however, the license clearly prohibits AlphaGenome use to train machine learning models.

From a technical point of view, eQIF comes with a small computational burden, enabling genome-wide pre-computation of scores, which makes it easily accessible for computational and experimental studies. However, eQIF is based on the precomputed Enformer principal components (PCs) as an input, which is only available for common SNPs from the 1000G cohort (20). Thus, at its current state, extending it for rare variants would require running the resource-intensive Enformer model for a tremendous number of predictions first. Faster future deep-learning-based models could grant a foundation for integrating rare variants into eQIF.

### eQIF’s impact on prioritizing the gene-regulatory genome

After extensive evaluation of eQIF’s performance, we investigated its molecular properties. High-scoring variants tended to be enriched at evolutionary conserved sites, specifically located within TFBS footprints of enhancers and promoters. TFBS footprints represent transcription factor binding measured by open chromatin assays. Previous analysis indicated that they are both highly relevant in gene regulation and also hypermutated (37). Because of the high number of mutations, causal variant discovery within TFBS footprints is challenging. The presence of top-scoring eQIF variants positively correlated with TFBS information content. Together with the enrichment of high eQIF scores at evolutionary conserved footprint sites, the TFBS correlation indicates that eQIF can be integrated for causal variant prioritization within TFBS footprints and thus to significantly reduce the search space of the gene-regulatory genome in the future.

## Data availability

The top 20 Enformer PCs are available under a CC-BY 4.0 license at https://console.cloud.google.com/storage/browser/dm-enformer/variant-scores/1000-genomes/enformer/PC20;tab=objects?prefix=&forceOnObjectsSortingFiltering=false.

GTEx v. 8 fine-mapping eQTL credible set collection is available at https://storage.googleapis.com/adult-gtex/bulk-qtl/v8/fine-mapping-cis-eqtl/GTEx_v8_finemapping_CAVIAR.tar.

The CAUSALdb2 v. 2.1 credible set collection is available at https://onedrive.live.com/?id=%2Fpersonal%2Ff7bb08e3479fe7ea%2FDocuments%2FCASUALdb&listurl=%2Fpersonal%2Ff7bb08e3479fe7ea%2FDocuments&ithint=folder&e=PaGHKZ&migratedtospo=true&redeem=aHR0cHM6Ly8xZHJ2Lm1zL2YvcyFBdXJubjBmakNMdjNna3NKSXdVQnA3Q3F1UVJWP2U9UGFHSEta&ga=1.

CADD v. 1.7 precomputed scores are available at https://cadd.bihealth.org/download.

Precomputed LINSIGHT scores are available at http://compgen.cshl.edu/LINSIGHT/LINSIGHT.bw.

Precomputed DVAR scores are available at https://www.vumc.org/cgg/dvar.

Precomputed AVI scores are available at https://alphagenome.google/atlas after registration.

phyloP scores for vertebrate, mammal, and primate alignments are available at https://hgdownload.soe.ucsc.edu/goldenPath/hg38/phyloP100way/hg38.phyloP100way.bw, https://hgdownload.soe.ucsc.edu/goldenPath/hg38/phyloP447way/hg38.phyloP447way.bw, and https://hgdownload.soe.ucsc.edu/goldenPath/hg38/phyloP447way/hg38.phyloP447wayPrimates.bw, respectively.

The code and pre-trained model to run Sei predictions are available at https://github.com/FunctionLab/sei-framework

Gene-regulatory annotation from the SCREEN v. 3 database can be downloaded from https://downloads.wenglab.org/V3/GRCh38-cCREs.bed.

TFBS footprint annotation can be obtained from https://resources.altius.org/~jvierstra/projects/footprinting.2020/consensus.index/consensus_footprints_and_collapsed_motifs_hg38.bed.gz.

TFBS data is accessible on the JASPAR 2026 website https://jaspar.elixir.no/downloads/.

The MEME Suite including the AME and FIMO tools can be obtained from https://meme-suite.org/meme/.

The GENCODE v.44 dataset can be obtained from https://ftp.ebi.ac.uk/pub/databases/gencode/Gencode_human/release_44/gencode.v44.annotation.gtf.gz

The hg38 reference genome can be obtained from https://hgdownload.soe.ucsc.edu/goldenpath/hg38/bigZips/hg38.fa.gz

eQIF is available running under a CC-BY 4.0 license. Genome-wide predictions for all European SNPs are publicly available on Zenodo (DOI = 10.5281/zenodo.23733502). The source code to reproduce our results is available on GitHub: https://github.com/pancinf/eQIF.git.

## Acknowledgements

The graphical abstract was created using BioRender. ChatGPT was used to assist with coding, improve the clarity and readability of the manuscript. Furthermore, ChatGPT was used for critical review and revision of the manuscript. For grammar checking and correction we used Grammarly and ChatGPT. All AI-assisted modifications were carefully reviewed and verified by the authors, who take full responsibility for the accuracy, integrity, and content of the manuscript.

## Author contributions

A.W.S., S.O., and H.L. conceived and directed the study.

A.W.S. conducted model development and data analysis.

A.W.S., S.O., H.L., and J.R. drafted and revised the manuscript.

All authors approved the final manuscript and contributed critical revisions to its intellectual content.

