## Supplementary_Figures for "Unsupervised prioritization of gene-regulatory variants with isolation forests"

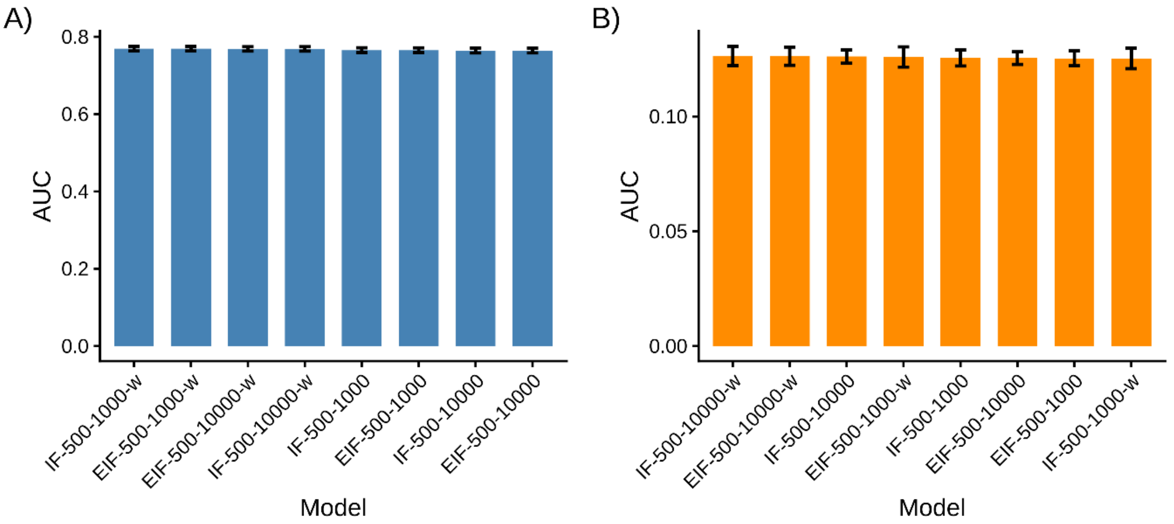


***Suppl. Fig 1: Parameter tuning of eQIF for eQTL classification.***
*A) shows the AUCs of ROC and B) the AUC of PRC analysis for eight isolation forest configurations. Hyperparameter optimization was performed using five repeated random 80:20 training-test splits (Monte Carlo cross-validation) of the complete genome-wide variant dataset. In each iteration, models were trained on 80% of the variants and evaluated on the remaining 20%, restricted to variants overlapping the predefined eQTL cohort variants. The parameters ndim and sample_size, as well as feature weighting according to PC rank, were evaluated, while ntrees was fixed at 500. Model names start either with "IF" in case of ndim = 1 or "EIF" in case of ndim = 2. The second and third model name entries describe the values set for ntrees and sample_size, respectively. "-w" indicates that the model used weighted feature selection. The 5 test sets contained 2536, 2499, 2527, 2475, 2523, case variants respectively, and 153401, 153013, 153823, 153658, and 153512 control variants, respectively.*

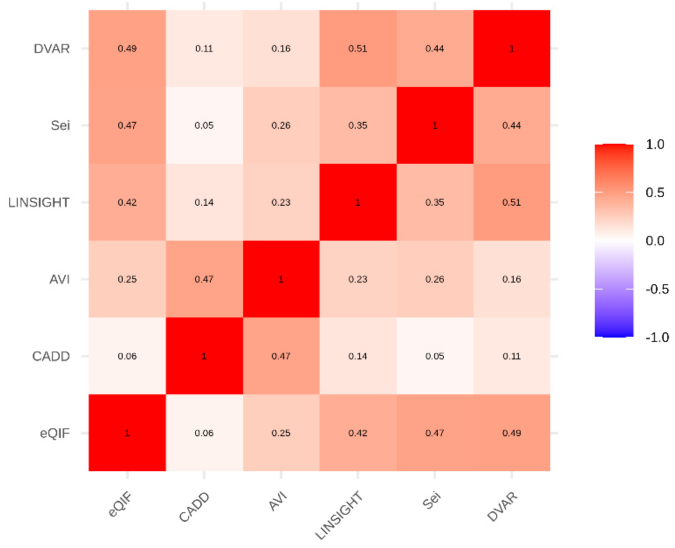


***Suppl. Fig 2: Correlation of variant impact prediction scores.***
*Spearman's rho was computed between all considered models for all variants within the eQTL cohort.*

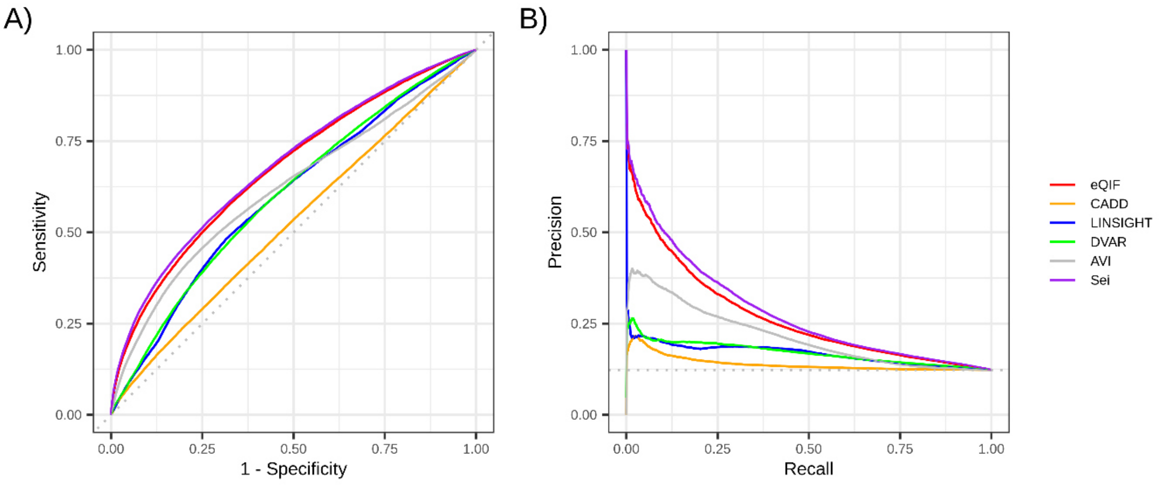


***Suppl. Fig 3: MPRA classification analysis.***
*A) shows the ROC and B) the PRC analysis. MPRA classification was evaluated on a positive group of variants fine-mapped eQTL or GWAS variants, which showed allelic differences in gene expression in MPRA analysis and a negative group of MPRA-negative variants. eQIF was compared to CADD v. 1.7, LINSIGHT, DVAR, AVI, and seqclass_max_absdiff from Sei. The number of case and control variants is 32,212 and 229,559, respectively.*

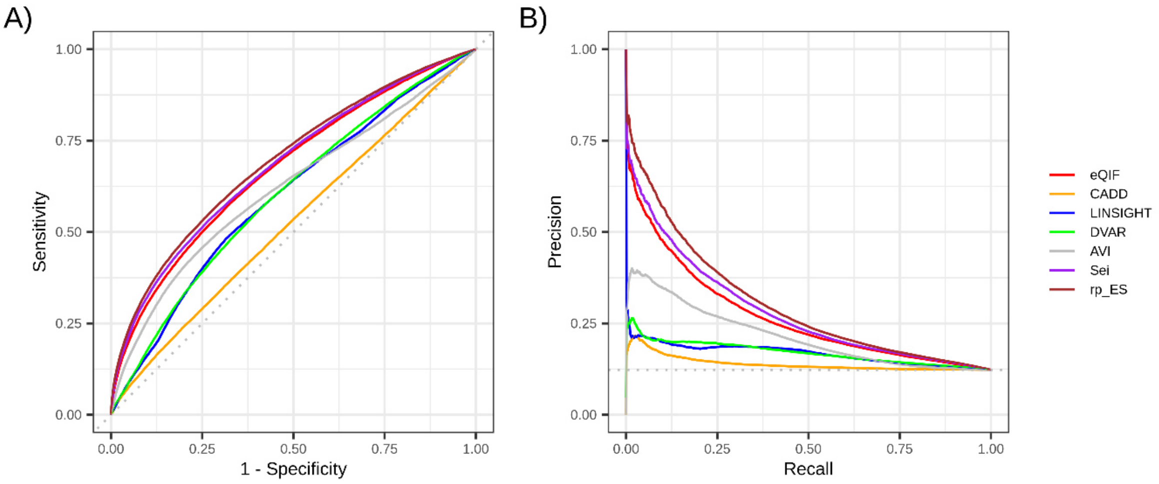


***Suppl. Fig 4: MPRA classification analysis including a rank product of eQIF and Sei.***
*A) shows the ROC and B) the PRC analysis. MPRA classification was evaluated on a positive group of variants fine-mapped eQTL or GWAS variants, which showed allelic differences in gene expression in MPRA analysis and a negative group of MPRA-negative variants. eQIF was compared to CADD v. 1.7, LINSIGHT, DVAR, AVI, seqclass_max_absdiff from Sei, and the rank product (rp_ES) of eQIF and seqclass_max_absdiff (computed across the entire dataset). The number of case and control variants is 32,212 and 229,559, respectively.*

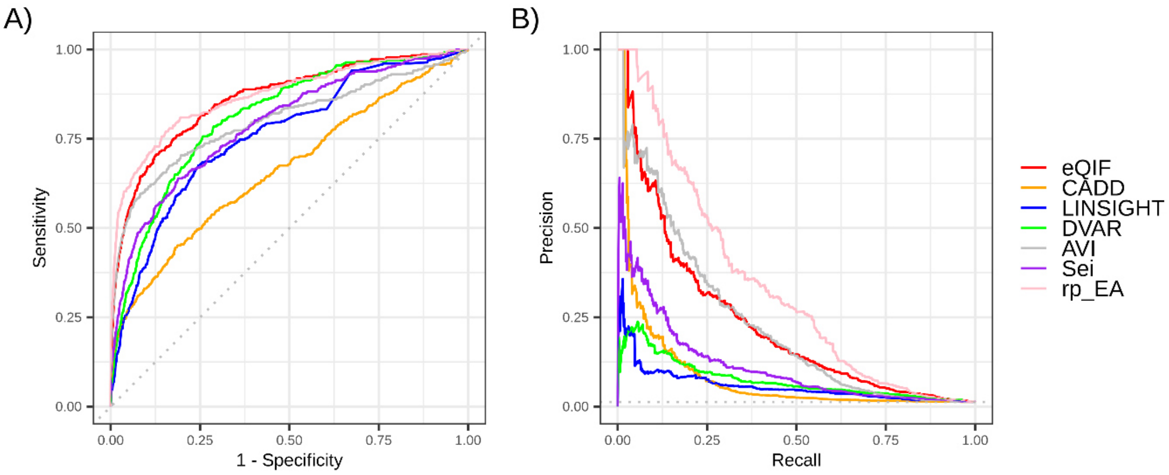


***Suppl. Fig 5: Comparing eQIF with previous tools and a rank product combination with AVI for gene-regulatory GWAS classification.***

*A) ROC and B) PRC analysis. GWAS-eQTL classification was evaluated on a positive group of variants strongly fine-mapped by both the CAUSALdb2 dataset and GTEx v.8 and a control dataset of variants with low fine-mapping probabilities. eQIF was compared to CADD v. 1.7, LINSIGHT, DVAR, AVI, seqclass_max_absdiff from Sei, and the rank product of eQIF and AVI (computed across the entire dataset). The number of case and control variants is 357 and 27,394, respectively.*

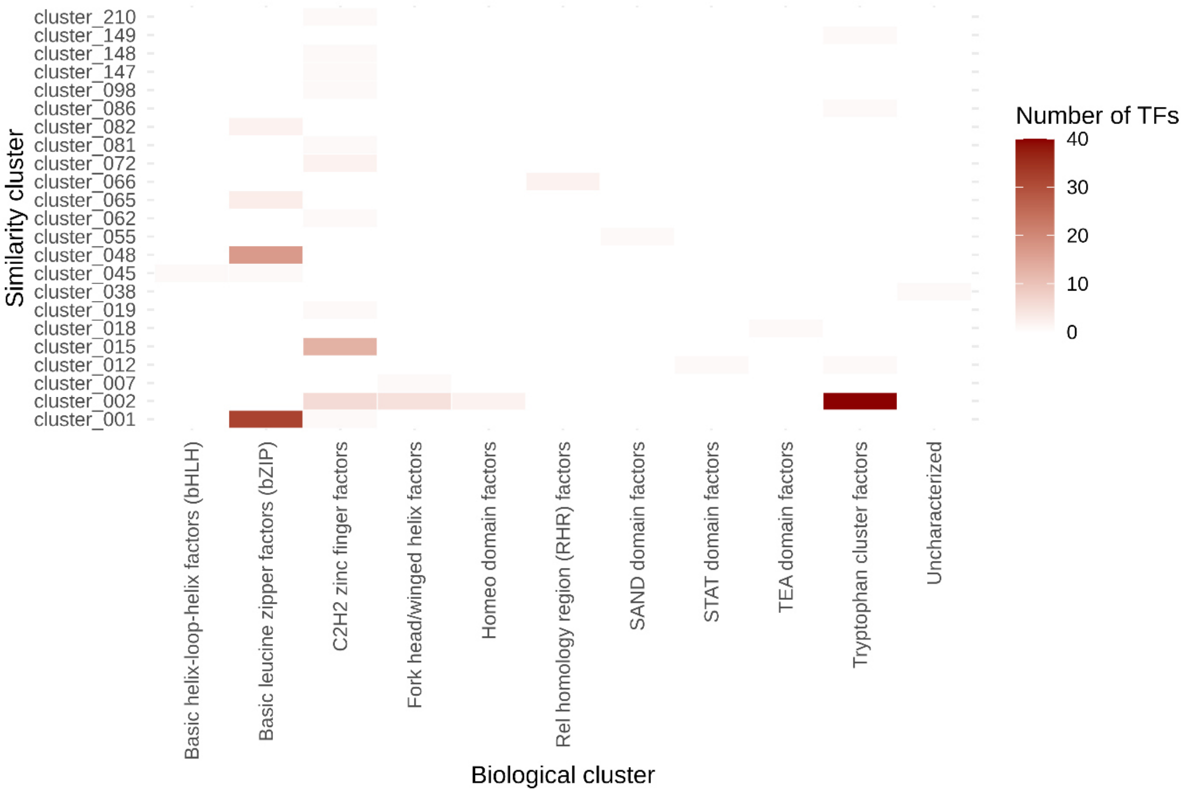


***Suppl. Fig 6: Clustering of TFBS enriched at the top 5 K eQIF SNPs.***

*142 TFBS motifs were discovered to be significantly enriched and of high fold change at top 5,000 eQIF SNPs. The TFBS were clustered by motif similarity (vertical annotation) and binding domain annotation of the corresponding protein (horizontal annotation). Clustering information was obtained from the Jaspar database.*
